# Integrative genomics analysis identifies *ACVR1B* as a candidate causal gene of emphysema distribution in non-alpha 1-antitrypsin deficient smokers

**DOI:** 10.1101/189100

**Authors:** Adel Boueiz, Robert Chase, Andrew Lamb, Sool Lee, Zun Zar Chi Naing, Michael H. Cho, Margaret M. Parker, Craig P. Hersh, James D. Crapo, Andrew B. Stergachis, Ruth Tal-Singer, Dawn L. DeMeo, Edwin K. Silverman, Peter J. Castaldi, for the COPDGene investigators

**Affiliations:** Channing Division of Network Medicine, Brigham and Women’s Hospital, Harvard Medical School, Boston, MA, 02115, USA; Pulmonary and Critical Care Medicine, Brigham and omen’s Hospital, Harvard Medical School, Boston, MA, 02115, USA; Pulmonary Medicine, National Jewish Health, Denver, CO, 80206, USA; Division of Genetics, Brigham and Women’s Hospital, Harvard Medical School, Boston, MA, 021 USA; GSK, King of Prussia, 19406, PA; General Medicine and Primary Care, Brigham and Women’s Hospital,Harvard Medical School, Boston, MA, 02115, USA.

**Author notes:** **Corresponding Author**: Peter J. Castaldi, MD, MSc, Channing Division of Network Medicine, Brigham and Women’s Hospital, 181 Longwood Avenue, Boston, MA, 02115. **Authors’ email addresses:** Adel Boueiz, Robert Chase, Andrew Lamb, Sool Lee, Zun Zar Chi Naing, Michael H. Cho, Margaret M. Parker, Craig P. Hersh, James D. Crapo, Andrew B. Stergachis, Ruth Tal-Singer, Dawn L. DeMeo, Edwin K. Silverman, Xiaobo Zhou, Peter J. Castaldi.

**Keywords:** Integrative genomics, Emphysema, Emphysema distribution, Chronic obstructive pulmonary disease

## Abstract

**Background:** Several genetic risk loci associated with emphysema apico-basal distribution (EABD) have been identified through genome-wide association studies (GWAS), but the biological functions of these variants are unknown. To characterize gene regulatory functions of EABD-associated variants, we integrated EABD GWAS results with 1) a multi-tissue panel of expression quantitative trait loci (eQTL) from subjects with COPD and the GTEx project and 2) epigenomic marks from 127 cell types in the Roadmap Epigenomics project. Functional validation was performed for a variant near *ACVR1B*.

**Results:** SNPs from 168 loci with P-values<5x10^-5^ in the largest GWAS meta-analysis of EABD *(Boueiz A. et al, AJRCCM 2017)* were analyzed. 54 loci overlapped eQTL regions from our multi-tissue panel, and 7 of these loci showed a high probability of harboring a single, shared GWAS and eQTL causal variant (colocalization posterior probability≥0.9). 17 cell types exhibited greater than expected overlap between EABD loci and DNase-I hypersensitive peaks, DNaseI hotspots, enhancer marks, or digital DNaseI footprints (permutation P-value < 0.05), with the strongest enrichment observed in CD4^+^, CD8^+^, and regulatory T cells. A region near *ACVR1B* demonstrated significant colocalization with a lung eQTL and overlapped DNase-I hypersensitive regions in multiple cell types, and reporter assays in human bronchial epithelial cells confirmed allele-specific regulatory activity for the lead variant, rs7962469.

**Conclusions:** Integrative analysis highlights candidate causal genes, regulatory variants, and cell types that may contribute to the pathogenesis of emphysema distribution. These findings will enable more accurate functional validation studies and better understanding of emphysema distribution biology.

## BACKGROUND

Chronic obstructive pulmonary disease (COPD) is a clinical syndrome with multiple, distinct clinical manifestations including the pathologic loss of lung tissue, i.e. emphysema. The phenotypic variability of COPD has prognostic and therapeutic implications [1-3]. Among patients with emphysema, patterns of lung destruction are often asymmetric [4], and this regional heterogeneity is an important factor in the severity of airflow limitation, disease progression, and the response to lung volume reduction procedures [5-9]. A recent genome-wide association study (GWAS) has identified five genomic loci associated with emphysema apico-basal distribution (EABD) in smokers without alpha-1 antitrypsin deficiency (4q13 near *SOWAHB*, 4q31 near *HHIP*, 8q24 near *TRAPPC9*, 10p12 near *KIAA1462,* and 15q25 near *CHRNA5*) [10]. These loci explain a small proportion of the estimated heritability of EABD, indicating that many more true associations have not yet been identified. In addition, the functional mechanisms of EABD causal variants have not yet been described.

GWAS variants tend to be located in regions of strong linkage disequilibrium (LD), making it difficult to identify the causal variant (or variants) in these regions from genetic association alone. In addition, GWAS-identified regions are frequently located in non-coding DNA and predominantly affect gene expression rather than directly affecting protein structure. This is supported by the observation that GWAS variants are enriched in regulatory domains, including enhancers and regions of open chromatin [11, 12]. We hypothesized that a number of EABD loci regulate gene expression and are located within genomic regions characterized by DNase-I hypersensitivity, enhancer activity, and transcription-factor binding. To test this hypothesis, we performed comprehensive fine-mapping of EABD-associated loci by integrating GWAS results with a large compendium of multi-tissue eQTL and cell-based epigenetic marks. This analysis identifies 7 high-confidence EABD-associated, gene regulatory loci in 42 tissues, and it implicates 17 cell types as likely to participate in EABD pathogenesis, with the strongest enrichment observed for T-cell subsets.

## RESULTS

### Characteristics of study participants included in the analysis

A previously published GWAS was performed using data from 6,215 non-Hispanic white and 2,955 African-American subjects from COPDGene, 1,538 subjects from ECLIPSE, and 824 subjects from GenKOLS with complete phenotype and genotype data *(Boueiz A. et al, AJRCCM 2017)*. The characteristics of these 11,532 subjects are shown in Table 1S, and the characteristics of the 385 COPDGene NHW subjects with available RNA-seq eQTL data are shown in Table 2S.

### Emphysema distribution GWAS signals colocalize with eQTLs

From the whole blood eQTL of the 385 COPDGene subjects, significant associations at 10% FDR were identified for 745,067 unique *cis*-eQTL SNP genotypes associated with the expression level of 17,187 unique transcripts (including protein coding genes, long non-coding RNA, and antisense transcripts) (Table 3S). These eQTLs were analyzed along with the multi-tissue GTEx eQTL data for a total number of unique *cis*-eQTL SNPs per tissue ranging between 129,647 (uterus) and 1,351,125 (thyroid) and unique eQTL transcripts per tissue ranging between 6,452 (uterus) and 27,618 (testis). The workflow of the GWAS-eQTL integration analysis and results are summarized in Figure 1.

**Figure 1.**
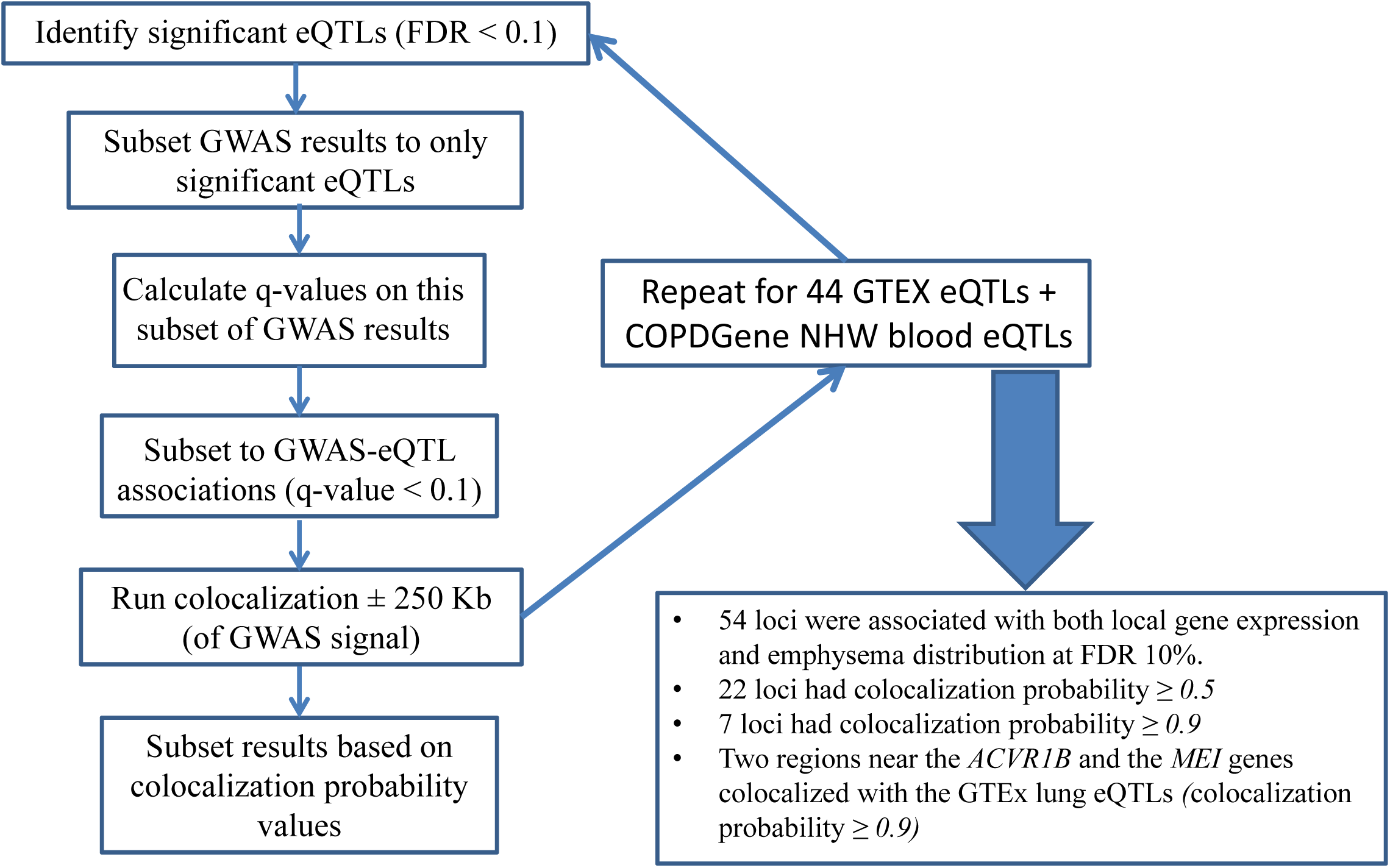
Workflow and summary of the results of the colocalization analysis of emphysema distribution-associated genetic variants with multi-tissue expression quantitative trait loci (eQTL) data from the Genotype-Tissue Expression (GTEx) project and whole blood eQTL from the COPDGene study.

Fifty-four genomic loci were associated with both local gene expression and emphysema distribution at FDR 10%, representing 32.1% (54/168) of the total number of candidate EABD-associated loci (GWAS P-value < 5x10^-5^). Given that overlap of GWAS and eQTL signals may occur by chance, a Bayesian test for colocalization was performed to distinguish overlap due to shared causal variants versus chance overlap. 13.1% (22/168) of candidate EABD loci had a reasonable likelihood (>50%) of harboring a shared causal variant for emphysema distribution and eQTL, and 4.2% of loci had a high likelihood (>90%) of harboring a shared causal variant (Table 1), including two regions that harbored genetic variants affecting the expression of *ACVR1B* and *MEI* in lung tissue (Figure 2). The complete set of colocalization results can be viewed interactively at https://cdnm.shinyapps.io/eabd_eqtlcolocalization/.

**Figure 2.**
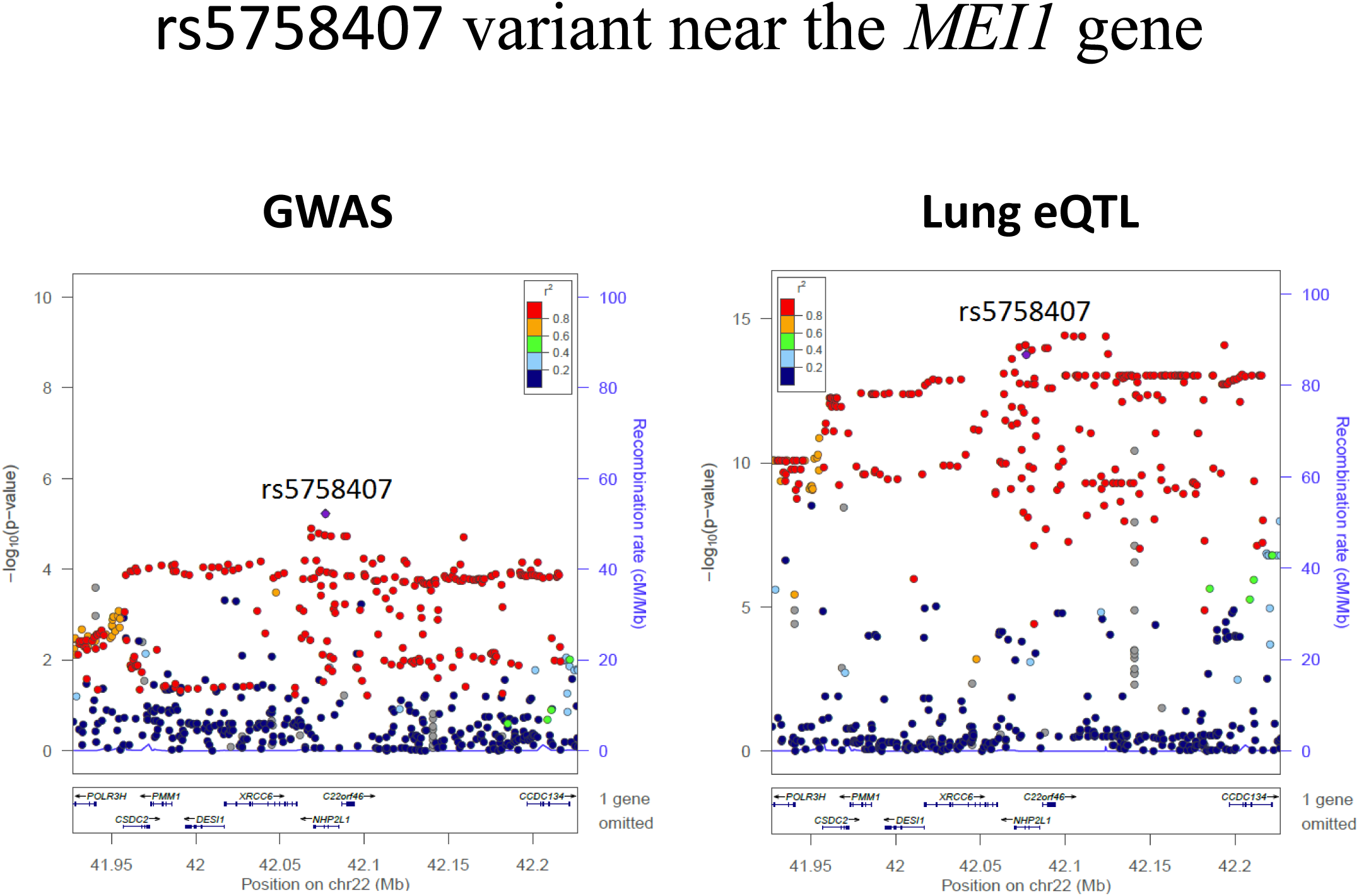
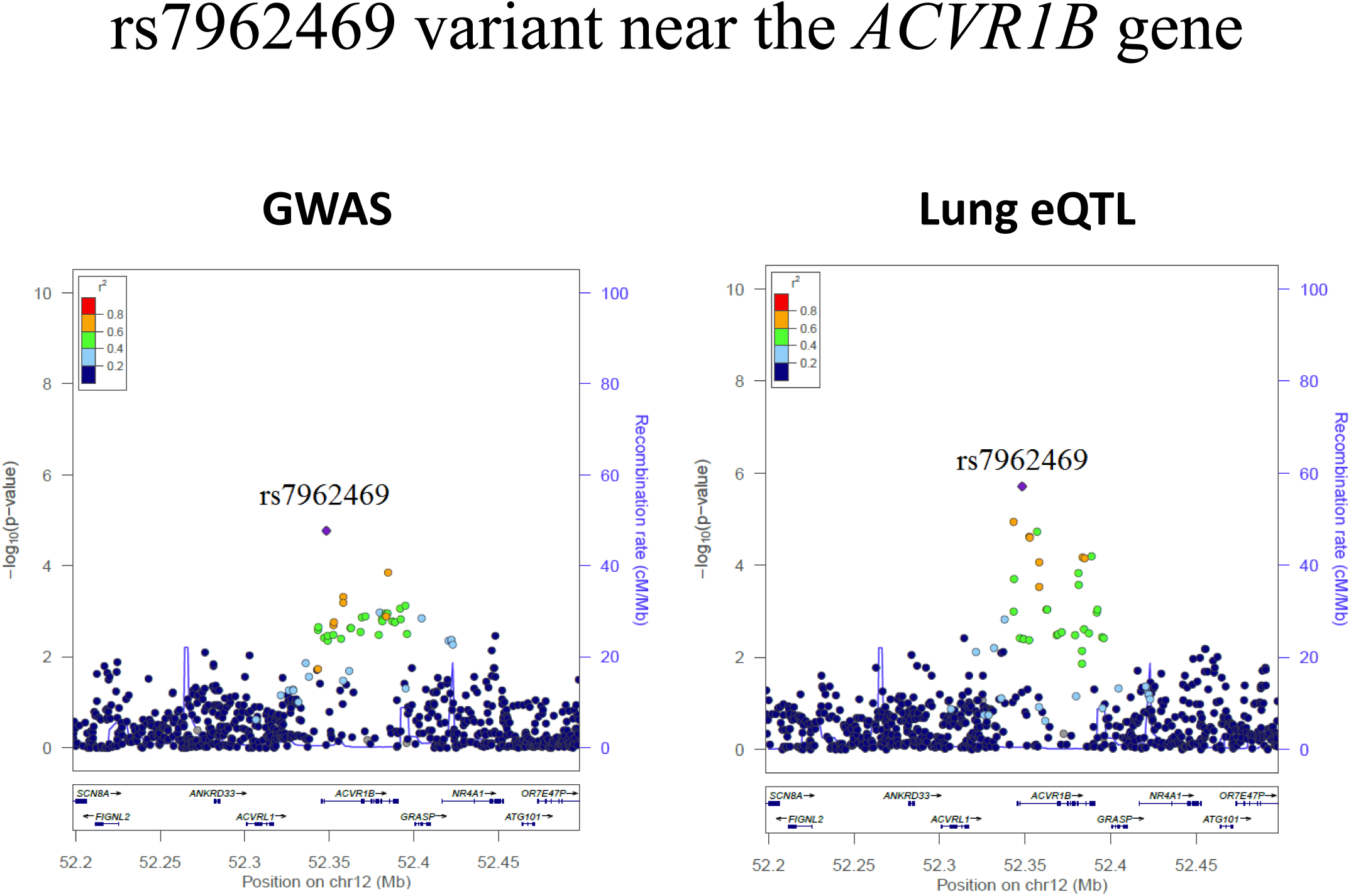
Genome-wide association study (GWAS) and lung expression quantitative trait loci (eQTL) locus plots for rs5758407 variant near the *MEI1* gene (Panel A) and rs7962469 variant near the *ACVR1B* gene (Panel B).

**Table 1.** Significant colocalizations of emphysema distribution-associated GWAS variants with eQTL from GTEx and the COPDGene study (colocalization probability ≥ 0.9) ordered by the lead SNP GWAS P-values.

| Lead SNP | Position | HUGO gene annotation | GWAS P-value | eQTL tissue | eQTL Q-value | Colocalization probability |
| --- | --- | --- | --- | --- | --- | --- |
| rs12914385 | 15:78898723 | *CHRNA5* | $1.70 \times 10^{-17}$ | Testis | 0.001 | 0.91 |
| rs2645694 | 4:77833947 | *SOWAHB* | $2.40 \times 10^{-8}$ | Skin: Sun-exposed (Lower leg) | 0.007 | 0.94 |
|  |  |  |  | Esophagus: Mucosa | 0.003 | 0.94 |
| rs35500465 | 19:1158884 | *SBNO2* | $1.80 \times 10^{-6}$ | Esophagus: Mucosa | 0.001 | 0.94 |
| rs5758407 | 22:42076956 | *MEI1* | $5.90 \times 10^{-6}$ | Cells: Transformed fibroblasts | $4.47 \times 10^{-18}$ | 0.93 |
| | | | | Lung | $1.59 \times 10^{-11}$ | 0.91 |
| | | | | Adipose: Subcutaneous | $1.53 \times 10^{-9}$ | 0.90 |
| rs17471079 | 13:99168721 | *STK24* | $1.10 \times 10^{-5}$ | Artery: Tibial | $1.47 \times 10^{-6}$ | 0.96 |
| rs7962469 | 12:52348259 | *ACVR1B* | $1.70 \times 10^{-5}$ | Lung | 0.0005 | 0.91 |
| rs4468504 | 14:107122186 | *IGHV3-66* | $2.70 \times 10^{-5}$ | Colon: Transverse | $6.45 \times 10^{-13}$ | 0.91 |
| Listed are all instances where the Bayesian colocalization results indicate that there is high posterior probability ($PP_4 \geq 0.9$) that the same causal variant is associated with emphysema distribution GWAS and eQTL signals. GWAS: Genome-wide association study; eQTL: Expression quantitative trait loci; Lead SNP: SNP with the lowest P-value for association in a 250-kb window; Colocalization probability: Posterior probability that the same causal variant is associated with emphysema distribution GWAS and eQTL signals. | | | | | | |

### Emphysema distribution-associated GWAS loci are enriched in T-cell subsets

To quantify the overlap between EABD-associated loci and epigenomic marks in cell types, we investigated whether candidate EABD-associated loci are enriched in gene regulatory regions identified by large scale functional studies performed by the ENCODE and Roadmap Epigenomics consortia [13, 14]. The four types of epigenomic marks that we studied span 39 to 127 diverse cell types and cover on average 0.44% to 2.87% of the genome per cell type (Table 4S). Figure 3 illustrates the workflow for the integrative analysis of GWAS and epigenomic marks.

**Figure 3.**
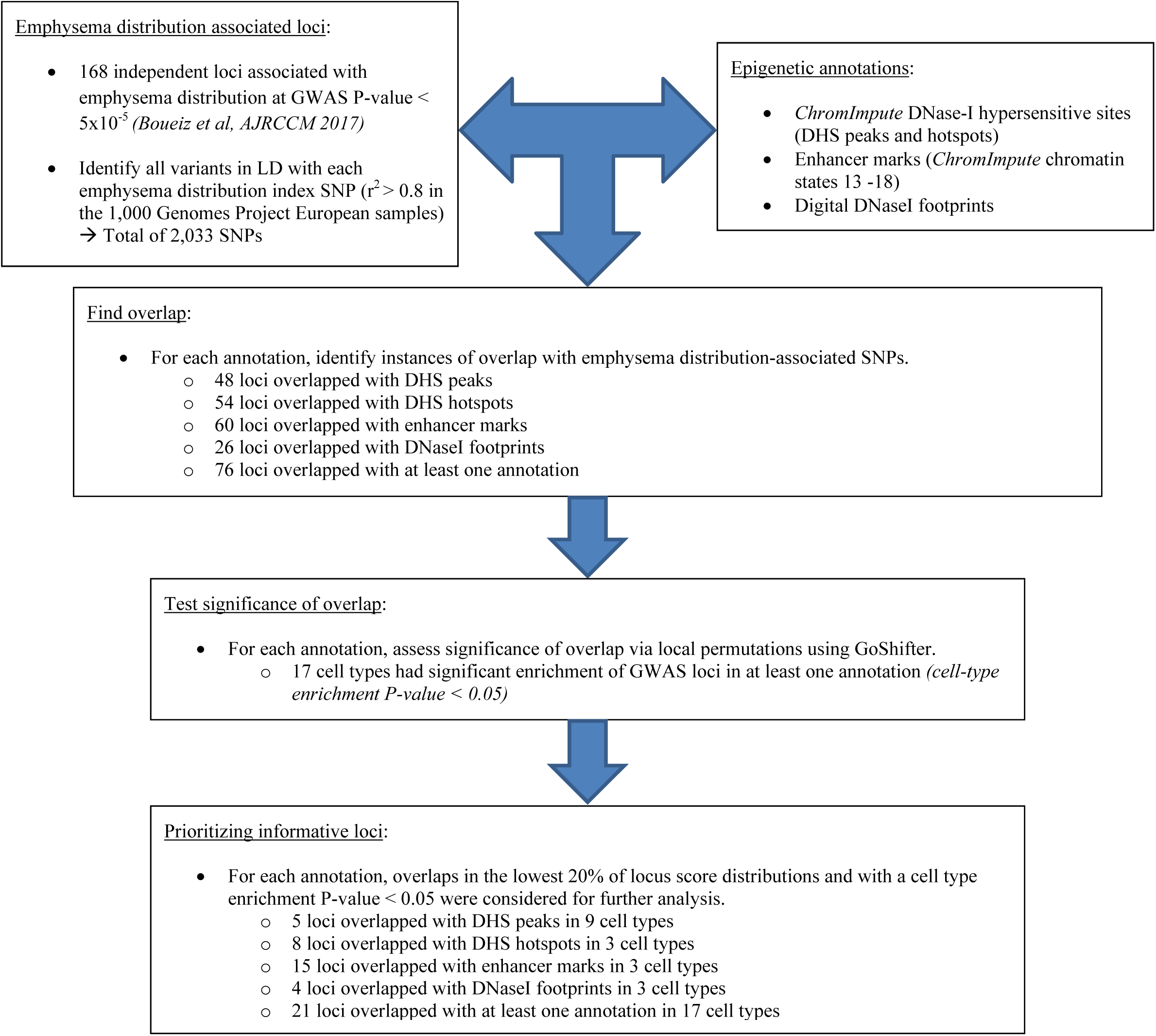
Workflow and summary of the integrative analysis of emphysema distribution-associated genetic variants with DNase-I hypersensitive peaks, DNaseI hotspots, enhancer marks, and digital DNaseI footprints.

Forty-five percent of EABD loci (76/168) overlapped at least one of the four studied epigenomic annotations in at least one cell type. Among those, 48 loci overlapped DHS peaks, 54 overlapped DHS hotspots, 60 overlapped enhancer marks, and 26 overlapped DNaseI footprints (28.6%, 32.1%, 35.7%, and 15.5% of EABD loci, respectively).

As with eQTL, some proportion of overlap between EABD loci and regulatory annotations may be due to chance rather than a causal link between the regulatory activity of a locus and its association to emphysema distribution. To better distinguish chance versus causal overlaps, we applied a previously published permutation approach that provides an estimate of the overall enrichment of GWAS loci in regulatory regions within a given cell type as well as a locus-specific score [15]. As illustrated in Figure 4, a total of 17 different cell types exhibited evidence of enrichment of EABD loci (Permutation p-value <0.05) in at least one set of epigenomic marks, with the most significant enrichment observed for CD4^+^, CD8^+^, and regulatory T cells. Prioritizing loci by cell type enrichment P-values < 0.05 and overlap locus scores in the lowest 20% of the overall locus score distribution for each epigenomic mark identified, 21 loci (12.5% of the EABD GWAS loci) in 17 different cell types overlapped at least one of the annotations (Table 2). The complete set of results can be viewed at https://cdnm.shinyapps.io/eabd_gwas_roadmap_goshifter/.

**Figure 4.**
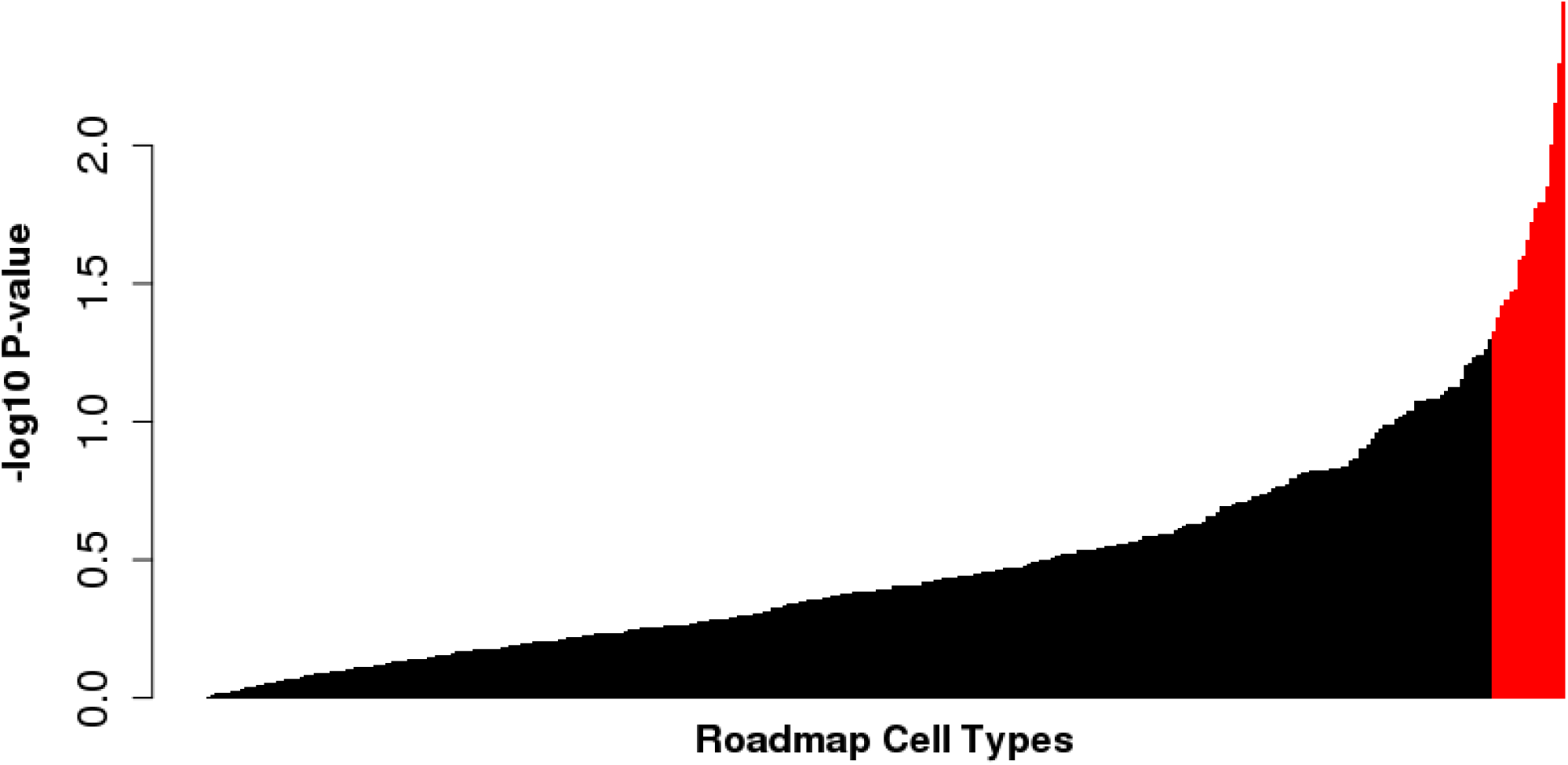
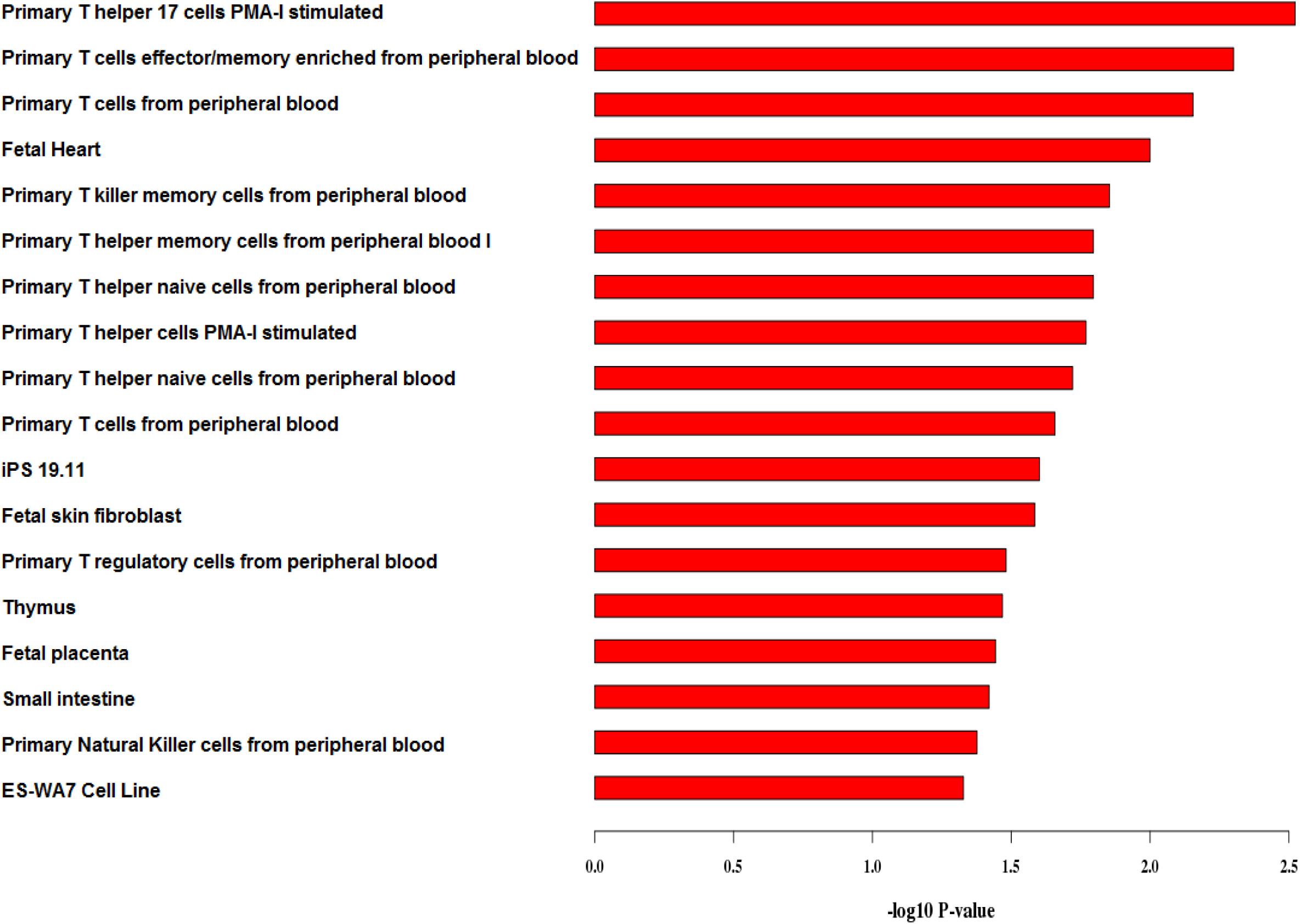
Plot of the cell type enrichment −log_10_P-value from the Genomic Annotation Shifter (GoShifter) analyses of the overlaps between emphysema distribution-associated genetic variants with DNase-I hypersensitive peaks, DNaseI hotspots, enhancer marks, and digital DNaseI footprints. The cell types with enrichment P-values less than 0.05 are highlighted in red in Figure 4A and are shown in more detail in Figure 4B.

**Table 2.**
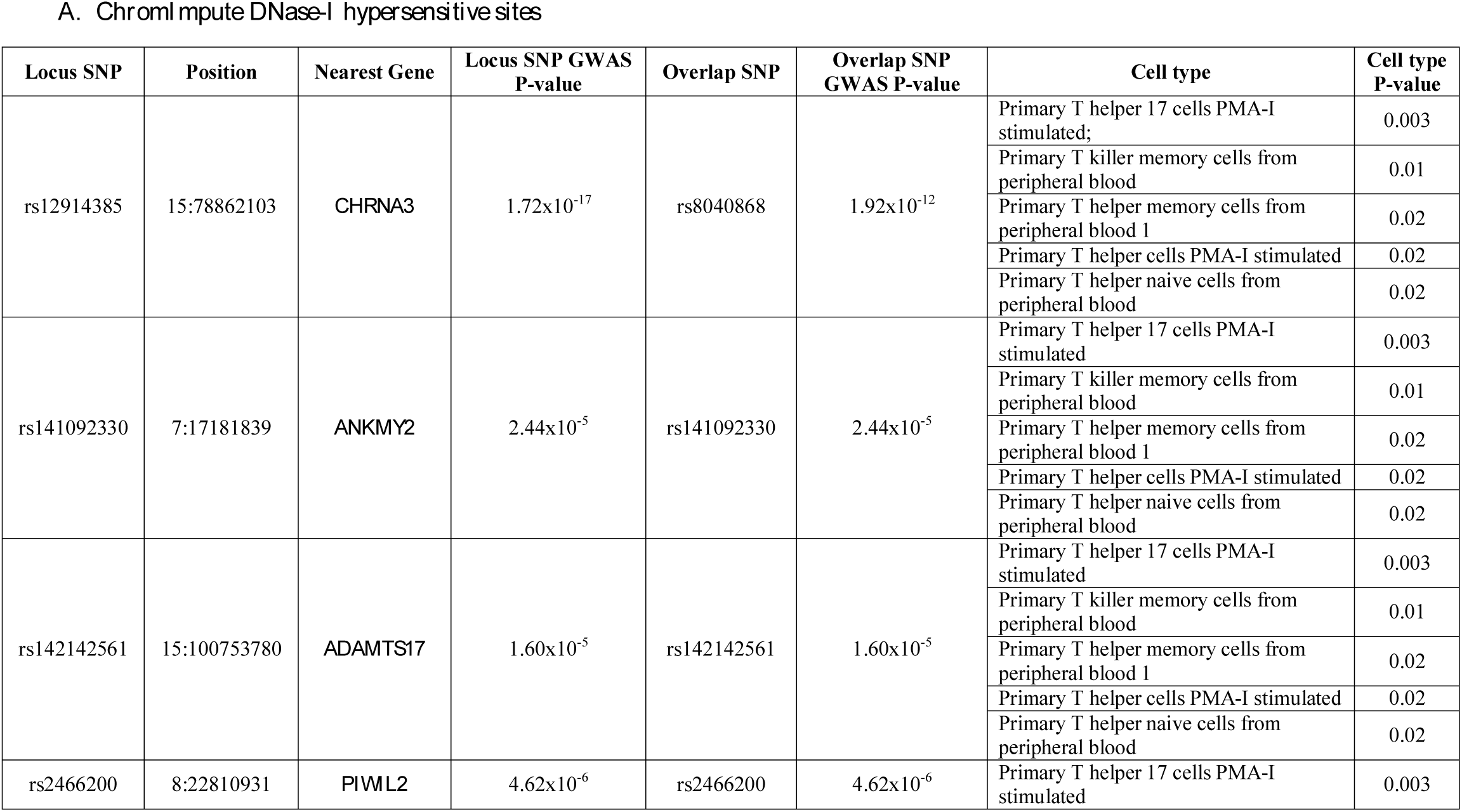
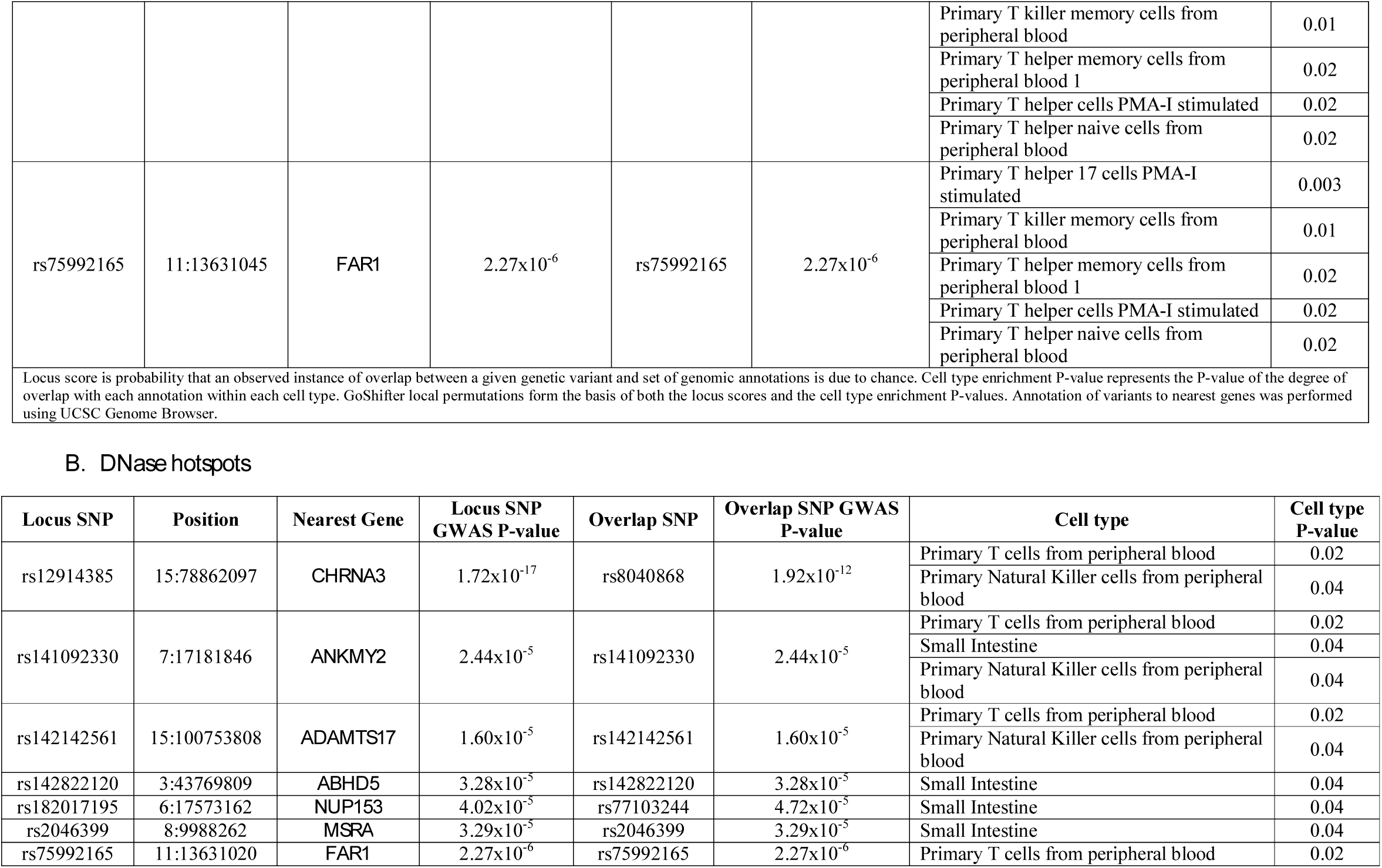
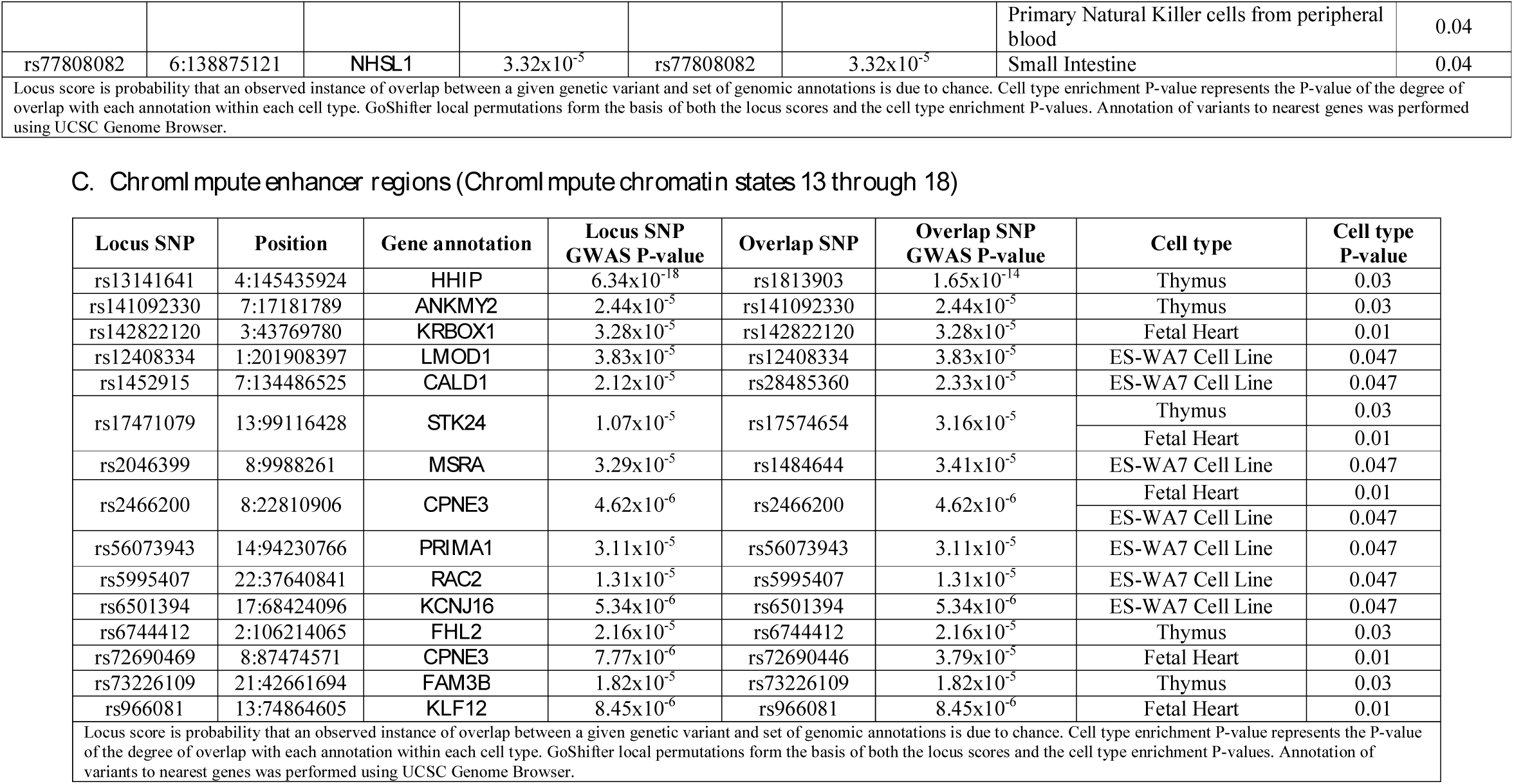
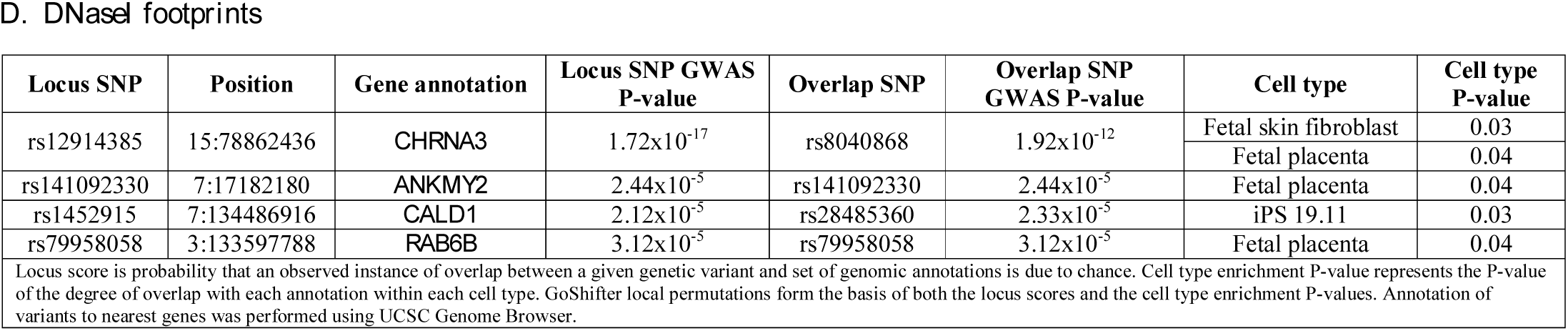
Significant overlaps between emphysema distribution-associated variants and epigenomic annotations (in the lowest 20% of locus score distributions and with a cell type enrichment P-value < 0.05).

### Overlaps between EABD GWAS loci, eQTL, and epigenomic marks

Forty-four percent of EABD GWAS-eQTL loci (24/54) also overlapped at least one of the epigenomic marks in at least one cell type (Table 3). Two of these loci (rs12914385 on chromosome 15 *(CHRNA5)* and rs17471079 on chromosome 13 *(STK24)*) had a GWAS-eQTL colocalization P-value ≥ 0.5, cell type epigenomic enrichment P-values < 0.05, and high priority locus scores.

**Table 3.** Number of overlap loci between emphysema apico-basal distribution GWAS, eQTL, and epigenomic marks.

|  |  | GoShifter |  |  |
| --- | --- | --- | --- | --- |
|  |  | Loci overlapping any annotation (76 loci) | Loci overlapping any annotation with cell type P-value < 0.05 (40 loci) | Loci overlapping any annotation with cell type P-value < 0.05 and in the lowest 20% of locus score distributions (21 loci) |
| eQTL | All GWAS-eQTL overlap loci (54 loci) | 24 | 15 | 6 |
| | GWAS-eQTL overlap loci with colocalization probability $\geq 0.5$ (22 loci) | 11 | 7 | 2 |
| | GWAS-eQTL overlap loci with colocalization probability $\geq 0.9$ (7 loci) | 5 | 4 | 2 |

### Functional validation of the rs7962469 variant near ACVR1B

Assuming that GWAS regions that colocalize with lung eQTLs may be more likely to play a causal role in emphysema distribution, we focused on the regions near the *ACVR1B* and the *MEI* genes for further functional prioritization. The pattern of association near *MEI* was nearly linear over a broad genomic region of high LD, making the identification of a single causal variant challenging. This, in combination with evidence of colocalization of rs7962469 with DNaseI hypersensitive regions in multiple cell types, led us to prioritize the region near *ACVR1B* for functional characterization. We used the PICS method to identify the 95% credible set of SNPs responsible for the GWAS association near *ACVR1B*, and the causal probability for rs7962469 was estimated to be 96%. Based on these results, we tested for an allelic expression effect of rs7962469 in human bronchial epithelial (16HBE) cells, and we observed that the G variant of rs7962469 has a significantly decreased expression relative to the A variant (Figure 5). The rs7962469 G variant has previously been shown to be associated with COPD susceptibility (OR: 1.13, SE: 1.03; GWAS P-value: 0.002) [16] and with upper lobe emphysema predominance (effect size: 0.06, SE: 0.01, GWAS P-value: 1.7x10^-5^) [10].

**Figure 5.**
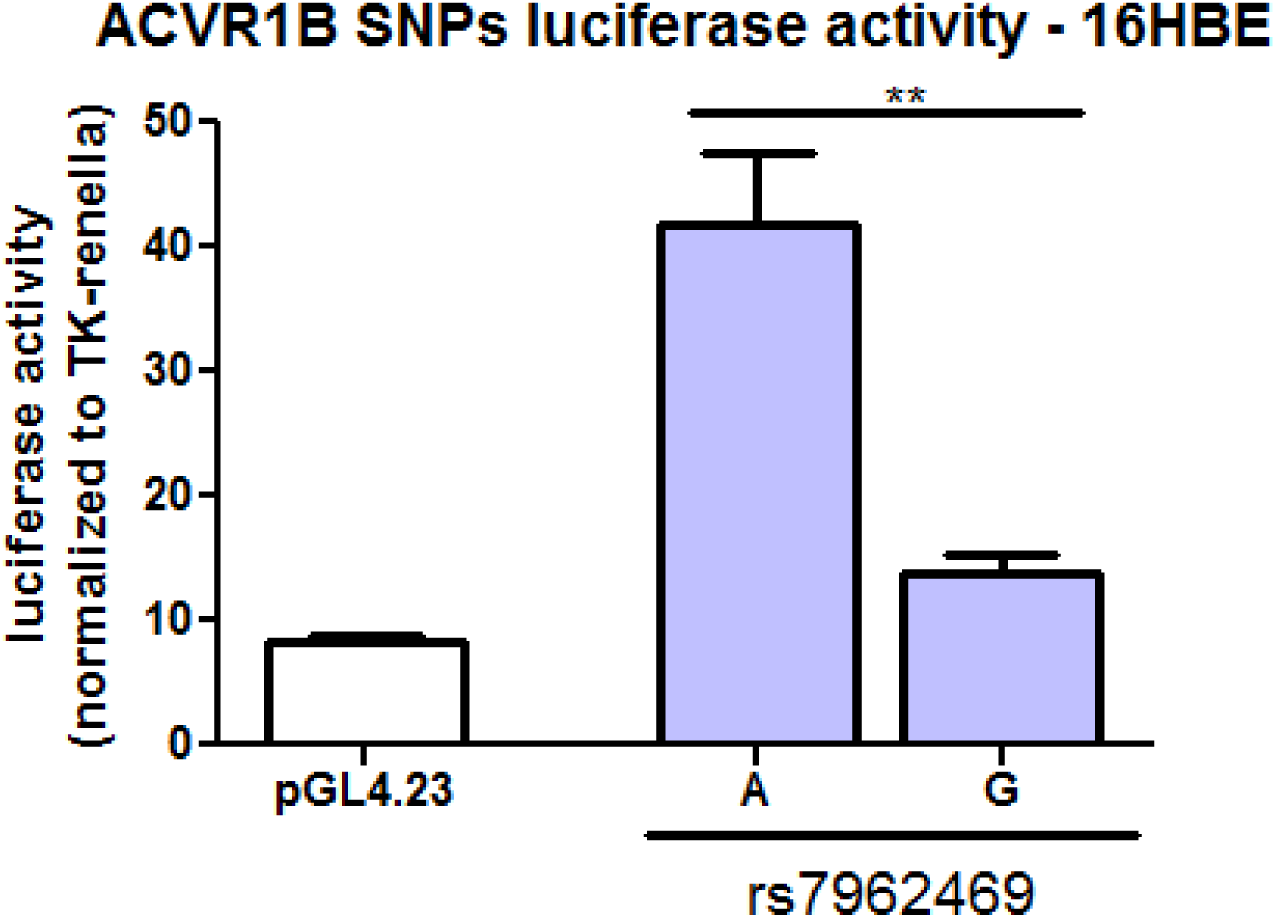
Reporter assays of the rs7962469 variant near the *ACVR1B* gene in Human Bronchial Epithelial (16HBE) cells showing significantly increased expression of the A variant relative to the G variant *(P-value: 0.008)* and the empty Luciferase vector (pGL4.23) *(P-value: 0.00002)*. The P-value comparing the G variant to pGL4.23 is 0.02. **: P-value <0.05.

## DISCUSSION

Using a multi-cohort GWAS of EABD, multi-tissue eQTL from 45 tissues, and epigenomic marks from 127 cell types, we performed an integrated genetic-epigenomic study to further our functional understanding of common variants associated with this specific manifestation of COPD. eQTL colocalization analysis identified strong evidence of a shared causal variant responsible for observed GWAS and eQTL associations in 7 distinct loci in multiple tissues. GWAS-epigenomic mark enrichment was observed for 17 cell types, with the strongest enrichment in CD4^+^, CD8^+^, and regulatory T cells. A region near the promoter of the *ACVR1B* gene demonstrated strong colocalization in lung eQTL data and lies within a DNase-I hypersensitive region that is active in multiple cell types. Reporter assays confirmed allele-specific regulatory activity for the EABD-associated variant, rs7962469, near *ACVR1B* with the G allele associated with decreased reporter gene expression, increased COPD susceptibility, and upper lobe emphysema predominance.

Efforts to understand how genetic variation contributes to common diseases increasingly focus on the regulation of gene expression [17-19]. The enrichment of *cis*-eQTL SNPs has been demonstrated for GWAS-identified loci in COPD [20], and EABD GWAS-eQTL overlap has been previously reported for EABD [21]. With the use of RNA-seq in multiple tissues in GTEx and in blood from subjects with COPD from the COPDGene study, the current study confirmed the previously reported EABD GWAS-eQTL overlap at the 15q25 locus with formal colocalization testing and added evidence for genetic control of gene expression at 12 other emphysema distribution-associated loci.

GWAS-identified loci for some phenotypes are enriched for regulatory marks in tissues that are relevant to the phenotype [21-25]. In the case of emphysema distribution, we observed GWAS-regulatory region overlap in multiple tissue and cell types, highlighting the biological complexity of emphysema and the fact that a significant amount of the regulatory genome is active across multiple tissues and cell types. While some disease loci lie within cell type-specific regulatory regions [26], our results are consistent with Boyle et al.’s work that provided evidence that many complex traits are driven by regulatory variants that tend to be active across many cell types and tissues [27].

Because most GWAS loci discovered to date lie outside of coding regions, it is likely that these loci affect gene regulation. 32.1% of the EABD-associated GWAS loci overlap with eQTL and 45.2% overlap with epigenomic annotations. However, our analysis indicates that a significant proportion of these overlaps may be due to chance, because only 13.1% of candidate EABD regions overlapped an eQTL with a colocalization probability ≥ 0.5, and only 12.5% overlapped an epigenomic mark with a high priority locus score. Furthermore, only 1.2% (2 loci) had a GWAS-eQTL colocalization P-value ≥ 0.5, cell type epigenomic enrichment P-values < 0.05, and high priority locus scores. Our observation that a notable proportion of observed GWAS overlaps with regulatory signals may be due to chance is consistent with previous observations in multiple sclerosis [28]. It is likely that this observation reflects the high prevalence of regulatory activity in the genome. However, the enrichment methods that we used suffer from important limitations. The Bayesian colocalization method assumes the presence of only one causal variant in a given locus for both GWAS and eQTL signals; this assumption reduces the accuracy of results when the locus contains multiple causal variants [17, 29]. The GoShifter method penalizes regions that are dense in epigenomic annotations and may therefore decrease the power of causal variant identification [15].

It is also interesting to consider the proportion of GWAS signals for which no regulatory overlaps were identified. These instances of non-overlap could be due to false positive GWAS associations at the reduced stringency levels used for this analysis, limited power of the included eQTL analyses [17], limited assessment of the dynamic nature of epigenomic marks in included cell type data, and lack of representation of relevant tissues and cell types for emphysema in GTEx and the Roadmap Epigenomics project, respectively. It is also possible that EABD causal variants affect aspects of gene regulation not observed in our data, such as splicing or post-transcriptional regulation [17].

It has been recognized for many years that emphysema often has a predilection for the upper lobes and subpleural areas in non-alpha-1 antitrypsin deficient smokers [10, 30]. The cause for these regional differences in emphysema distribution is not well understood but has been attributed to regional differences in perfusion, transit time of leukocytes, clearance of deposited dust, mechanical stress and pleural pressure [30-33]. This study provides compelling evidence to support a causal role for the adaptive immune response in EABD, with the strongest enrichment of EABD loci in regulatory activity observed in CD4^+^, CD8^+^, and regulatory T cells. Furthermore, based on the results of the genomics integrative analysis and the confirmatory reporter assay, our study prioritized the rs7962469 variant, near the *ACVR1B* gene, as a candidate causal variant in emphysema distribution.

*ACVR1B*, also known as *ALK-4*, acts as a transducer of activin-like ligands that are growth and differentiation factors belonging to the transforming growth factor-β (TGF-β) superfamily of signaling proteins. Although *ACVR1B* has not previously been associated with emphysema distribution, prior genetic studies have demonstrated an association of gene polymorphisms of the TGF-β superfamily with COPD [34, 35]. A genome-wide association meta-analysis of 3,497 subjects with severe COPD identified a genome-wide significant association with a locus previously reported to affect the gene expression of *TGFB2* [16]. In addition, *TGFB2* expression levels were reduced in a set of Lung Tissue Research Consortium COPD lung tissue samples compared with controls [16]. A network analysis incorporating COPD GWAS and protein-protein interaction (PPI) data included *ACVR1B* gene in a 10 gene consensus network module associated with COPD case-control status [36]. In addition, differential expression of *ACVR1B* has been found in the epithelial cells of a subset of smokers with lung cancer and in bone marrow micro-metastases from lung cancer patients [37, 38]. More work is warranted to elucidate the role of *ACVR1B* in COPD and emphysema.

The strengths of this study are 1) the use of comprehensive compendia of eQTL and epigenetic marks in multiple tissues and cell types for a novel COPD-related phenotype, 2) application of Bayesian and permutation-based methods to assess the significance of observed overlaps accounting for the genomic abundance of candidate eQTLs and epigenomic annotations, and 3) functional validation and demonstration of allele-specific enhancer activity for a candidate causal variant near *ACVR1B* prioritized by this integrative method.

This study also has important limitations. First, we limited our analysis to *cis*-eQTLs, excluding other classes of gene regulatory variants including *trans*-eQTLs, isoform ratio QTLs, and variants implicated by allele-specific expression analyses. Future studies will be strengthened by the inclusion of these emerging features of eQTL studies [39, 40]. Second, the colocalization and enrichment methods that we used have limitations. The colocalization method does not account for multiple independent signals, and the GoShifter approach is biased against regions that are dense in epigenomic annotations and may therefore decrease the power of causal variant identification [15, 17, 29].

## CONCLUSION

This study provides proof of concept for the effectiveness of an approach to leverage multi-tissue compendia of eQTLs and multi-cell compendia of epigenetic marks to refine and characterize functional regulatory GWAS loci. Enrichment analyses implicated a wide range of cells and tissues, emphasizing the importance of having comprehensive compendia of regulatory annotation with respect to tissues, cell types, diseases, and environmental exposures. Based on these integrative analyses, we prioritized and functionally validated a COPD and emphysema-associated variant involved in TGF-β signaling.

## MATERIALS AND METHODS

### Study populations

We analyzed 11,532 non-alpha-1 antitrypsin deficient current and former smokers with complete genotype and CT densitometry data from four cohorts: The Genetic Epidemiology of COPD study non-Hispanic whites (COPDGene NHW), COPDGene African-Americans (COPDGene AA), the Genetics of Chronic Obstructive Lung Disease (GenKOLS) and the Evaluation of COPD Longitudinally to Identify Predictive Surrogate Endpoints study (ECLIPSE). Detailed descriptions including study populations, genotyping quality control and genotyping imputation have been previously published [10, 41].

### CT measurements

Quantitative assessment of emphysema was performed using 3D SLICER density mask analysis (www.chestimagingplatform.org) to determine the percentage of lung voxels with attenuation lower than -950 Hounsfield units (%LAA-950) at maximal inspiration [42]. From these measurements, two correlated but complementary measures of emphysema distribution were constructed: 1) the difference between upper third and lower third emphysema *(diff950)* and 2) the ratio of upper third to lower third emphysema *(ratio950)* [10]. A rank-based inverse normal transformation was applied to both phenotypes to reduce the impact of outliers and deviations from normality [10]. In this current study, given that *ratio950* had a higher heritability and was associated with more genome-wide significant signals compared to *diff950* [10], we performed fine mapping of *ratio950*-associated variants.

### Peripheral blood gene expression

385 NHW subjects from the COPDGene study with completed peripheral blood RNA sequencing (RNA-seq) data were analyzed. All samples had RNA integrity number > 7 and RNA concentration ≥ 25 µg/µl (COPDGene Phase I dataset, January 6, 2017).

*RNA isolation and quality control:* Total RNA was extracted from PAXgene Blood RNA tubes using the Qiagen PreAnalytiX PAXgene Blood miRNA Kit. The extraction protocol was performed either manually or with the Qiagen QIAcube extraction robot.

*cDNA library preparation and sequencing:* Globin reduction and cDNA library preparation for total RNA was performed with the Illumina TruSeq Stranded Total RNA with Ribo-Zero Globin kit. Libraries were pooled and 75bp paired end reads were generated on the Illumina HiSeq2000 platform. Samples were sequenced to an average depth of 20 million reads.

*Read alignment, expression quantification, and sequencing quality control:* Reads were trimmed of the Truseq adapters using Skewer [43] with default parameters. Trimmed reads were aligned to GRCH38 genome using STAR [44]. Gene level counts were generated using RSubreads with the Ensemble GTF (version 81). Quality control was performed using the Fastqc [45] and RNA-SeQC [46] programs. Samples were included for subsequent analysis if they had >10 million reads, >80% of reads mapped, XIST and Y chromosome expression consistent with reported gender, <10% of R1 reads in the sense orientation, Pearson correlation ≥ 0.9 with other samples in the same library prep batch, and concordant genotype calls between RNA reads and DNA genotyping.

*cis-eQTL analysis:* Transcript-level expression count data were normalized for library size using the trimmed mean of M values method and then inverse normal transformed [47]. eQTL associations were tested for bi-allelic, autosomal SNPs with minor allele frequency (MAF) > 0.05 and mapping to a dbSNP 142 Reference SNP number. *cis-*eQTL analysis was performed for all SNPs within one megabase of the target gene using Matrix eQTL with a linear model adjusting for age, gender, library prep batch, 3 principal components of genetic ancestry, and 35 PEER factors of gene expression [48]. A total of 5,815,008 SNPs were tested for association with 27,277 transcripts. The threshold for significance was a false discovery rate (FDR) 10%, using the FDR procedure implemented in Matrix eQTL [49].

### eQTL-emphysema distribution GWAS colocalization analysis

We integrated emphysema distribution GWAS loci with multi-tissue eQTL data from the Genotype-Tissue Expression (GTEx) project *(version 6)* [10] (Download site: http://www.gtexportal.org/home/datasets; Date of download: November 18, 2015) and whole blood eQTL from 385 COPDGene NHW subjects. GWAS eQTL integration was performed as previously described [21]. Briefly, for each set of eQTL results, GWAS results were filtered to include only SNPs with a significant eQTL association at FDR 10%. In this reduced set of GWAS results, q-values were calculated using the procedure of Storey et al. for the GWAS P-values [50], and SNPs significant in both the eQTL and GWAS analyses at FDR 10% were retained. For each independent association, Bayesian colocalization tests were performed for all SNPs within a 250kb window of the lead GWAS variant at that locus to quantify the probability that the GWAS and eQTL associations were due to a single, shared causal variant [29]. This probability corresponds to the PP_4_ number described in the original publication. The workflow of this GWAS-eQTL integration analysis is summarized in Table 2.

### Enrichment in tissue-/cell type-specific chromatin states

Genetic variants associated with complex diseases have been shown to overlap regulatory enhancer elements [11, 12]. Based on this finding, we quantified the overlap between emphysema distribution GWAS SNPs (P-value < 5x10^-5^) and regulatory elements identified in the Roadmap Epigenomics project *(Release 9)* [51]. For the 127 Roadmap cell types, we downloaded *ChromImpute* imputed annotations that provide the most comprehensive human regulatory region annotation to date for large-scale experimental mapping of epigenomic information [52]. We analyzed *ChromImpute* DNase-I hypersensitive peaks and *ChromImpute* enhancer marks (defined as chromatin states 13 through 18) for all 127 cell types with available data. We also examined DNase-I hypersensitive hotspots that were available for 39 Roadmap cell types. Hotspots are broader regions of DNaseI hypersensitivity encompassing DNase peaks. The hotspot identification algorithm has been previously described [53]. We also analyzed digital DNaseI footprints (DGF) data from 42 uniformly processed cell types in Roadmap. DNaseI footprints are coverage troughs in deeply sequenced DNaseI hypersensitivity data that represent narrow genomic regions shielded from DNaseI digestion because of transcription factors, and thus represent likely transcription factor binding sites (TFBS).

DHS *ChromImpute* peaks were downloaded from http://egg2.wustl.edu/roadmap/data/byFileType/peaks/consolidatedImputed/narrowPeak/ on July 13, 2016. DHS hotspots were downloaded from http://egg2.wustl.edu/roadmap/data/byFileType/peaks/consolidated/broadPeak/DNase/ on February 20, 2015. Enhancer marks were downloaded from http://egg2.wustl.edu/roadmap/data/byFileType/chromhmmSegmentations/ChmmModels/imputed12marks/jointModel/final/ on Dec 23, 2015. DGF were downloaded from http://egg2.wustl.edu/roadmap/data/byDataType/dgfootprints/ on July 13, 2016.

To determine the extent of GWAS-epigenomic annotation overlap, we identified independent emphysema distribution GWAS signals at P-value < 5x10^-5^ within 1MB windows. We then used Genomic Annotation Shifter (GoShifter) to calculate the enrichment for these variants in Roadmap annotations (DHS, enhancer marks, DGF) [15]. This method uses a local permutation strategy to account for the local density of a given epigenomic mark and generates a score for each locus that can be used to prioritize loci where the overlap between a SNP and an annotation is particularly informative. 1,000 permutations were performed using LD information from the 1,000 Genomes EUR population with an r^2^ threshold of 0.8. Overlaps in the lowest 20% of locus score distributions and with a cell type enrichment P-value < 0.05 were considered for further analysis (Table 3).

### Shiny applications

Searchable tables of the formal metrics, results, and locuszoom plots for each genomic region was compiled and displayed using the R web framework shiny (http://shiny.rstudio.com/) and are available to the public as companion sites for this paper (https://cdnm.shinyapps.io/eabd_eqtlcolocalization/ *(GWAS-eQTL)* and https://cdnm.shinyapps.io/eabd_gwas_roadmap_goshifter/ *(GWAS-epigenomic annotations)).*

### Probabilistic Identification of Causal SNPs - rs7962469

The online Probabilistic Identification of Causal SNPs (PICS) algorithm is a fine mapping algorithm that calculates the probability that an individual SNP is a causal variant given the haplotype structure and observed pattern of association at the locus (https://pubs.broadinstitute.org/pubs/finemapping/pics.php) [54]. We used this algorithm to generate the 95% credible SNP set for the GWAS association identified in the region near the *ACVR1B* gene.

### Luciferase reporter assay - rs7962469

Two ∼500 base-pair long genomic segments including rs7962469 were obtained from Human Bronchial Epithelial (16HBE) cells heterozygous at rs7962469 and cloned into the sites of XhoI and BglII of a pGL4.23[luc2/minP] vector. Each luciferase construct was co-transfected with TK-Renilla, a luciferase control reporter, in 16HBE cells at ∼60-70% confluency by using Lipofectamine 3000 Reagent (Invitrogen), following the manufacturer’s protocol. Each luciferase construct was transfected in triplicate at a concentration of 300 ng per well and the TK-Renilla at 15 ng per well. Empty Luciferase vector, pGL4.23[luc2/minP], was also transfected in triplicates as a control. Promoter activity was quantified forty-eight hours post-transfection using the Dual-Luciferase Reporter Assay System (Promega) according to the manufacturer's protocol. Luminescence signals were captured in a Wallac VICTOR3 1420 plate reader (Perkin Elmer) and normalized by the Renilla luciferase readings for each well. The normalized values for each triplicate were then averaged. All plasmids used were confirmed by sequencing. Independent transfection and reporter assays were performed three times. Luciferase activity levels were assessed using Wilcoxon’s rank sum test; values were compared with a reference group within an experimental repeat, and results from multiple experiments were included. P-values less than were considered significant.

## DECLARATIONS

### Ethics approval and consent to participate

Informed consent was obtained from all study subjects and study approval was obtained from institutional review boards for all participating institutions.

### Availability of data and material

COPDGene data are available in the NCBI dbGaP database of genotypes and phenotypes under accession number phs000179.v1.p1.

### Consent for publication

Not applicable.

### Authors' contributions

Dr. Castaldi had full access to all of the data in the study, takes responsibility for the integrity of the data and the accuracy of the data analysis, had authority over manuscript preparation and the decision to submit the manuscript for publication.

*Study concept and design:* Boueiz, Castaldi

*Acquisition, analysis, or interpretation of data:* All authors

*Drafting of the manuscript:* Boueiz, Castaldi

*Critical revision of the manuscript for important intellectual content:* All authors

*Statistical analysis:* Boueiz, Castaldi *Obtained funding:* Castaldi, Crapo, Silverman

*Study supervision:* All authors

All authors gave final approval of the version to be published and have agreed to be accountable for all aspects of the work.

#### Acknowledgements

##### COPDGene Investigators – Core Units

*Administrative Core:* James Crapo, MD (PI), Edwin Silverman, MD, PhD (PI), Barry Make, MD, Elizabeth Regan, MD, PhD

*Genetic Analysis Core:* Terri Beaty, PhD, Nan Laird, PhD, Christoph Lange, PhD, Michael Cho, MD, Stephanie Santorico, PhD, John Hokanson, MPH, PhD, Dawn DeMeo, MD, MPH, Nadia Hansel, MD, MPH, Craig Hersh, MD, MPH, Peter Castaldi, MD, MSc, Merry-Lynn McDonald, PhD, Emily Wan, MD, Megan Hardin, MD, Jacqueline Hetmanski, MS, Margaret Parker, MS, Marilyn Foreman, MD, Brian Hobbs, MD, Adel Boueiz, MD, Peter Castaldi, MD, Megan Hardin, MD, Dandi Qiao, PhD, Elizabeth Regan, MD, Eitan Halper-Stromberg, Ferdouse Begum, Sungho Won, Sharon Lutz, PhD

*Imaging Core:* David A Lynch, MB, Harvey O Coxson, PhD, MeiLan K Han, MD, MS, MD, Eric A Hoffman, PhD, Stephen Humphries MS, Francine L Jacobson, MD, Philip F Judy, PhD, Ella A Kazerooni, MD, John D Newell, Jr., MD, Elizabeth Regan, MD, James C Ross, PhD, Raul San Jose Estepar, PhD, Berend C Stoel, PhD, Juerg Tschirren, PhD, Eva van Rikxoort, PhD, Bram van Ginneken, PhD, George Washko, MD, Carla G Wilson, MS, Mustafa Al Qaisi, MD, Teresa Gray, Alex Kluiber, Tanya Mann, Jered Sieren, Douglas Stinson, Joyce Schroeder, MD, Edwin Van Beek, MD, PhD

*PFT QA Core, Salt Lake City, UT:* Robert Jensen, PhD

*Data Coordinating Center and Biostatistics, National Jewish Health, Denver, CO:* Douglas Everett, PhD, Anna Faino, MS, Matt Strand, PhD, Carla Wilson, MS

*Epidemiology Core, University of Colorado Anschutz Medical Campus, Aurora, CO:* John E. Hokanson, MPH, PhD, Gregory Kinney, MPH, PhD, Sharon Lutz, PhD, Kendra Young PhD, Katherine Pratte, MSPH, Lindsey Duca, MS.

##### COPDGene Investigators – Clinical Centers

*Ann Arbor VA:* Jeffrey L. Curtis, MD, Carlos H. Martinez, MD, MPH, Perry G. Pernicano, MD

*Baylor College of Medicine, Houston, TX:* Nicola Hanania, MD, MS, Philip Alapat, MD, Venkata Bandi, MD, Mustafa Atik, MD, Aladin Boriek, PhD, Kalpatha Guntupalli, MD, Elizabeth Guy, MD, Amit Parulekar, MD, Arun Nachiappan, MD

*Brigham and Women’s Hospital, Boston, MA:* Dawn DeMeo, MD, MPH, Craig Hersh, MD, MPH, George Washko, MD, Francine Jacobson, MD, MPH

*Columbia University, New York, NY:* R. Graham Barr, MD, DrPH, Byron Thomashow, MD, John Austin, MD, Belinda D’Souza, MD, Gregory D.N. Pearson, MD, Anna Rozenshtein, MD, MPH, FACR

*Duke University Medical Center, Durham, NC:* Neil MacIntyre, Jr., MD, Lacey Washington, MD, H. Page McAdams, MD

*Health Partners Research Foundation, Minneapolis, MN:* Charlene McEvoy, MD, MPH, Joseph Tashjian, MD

*Johns Hopkins University, Baltimore, MD:* Robert Wise, MD, Nadia Hansel, MD, MPH, Robert Brown, MD, Karen Horton, MD, Nirupama Putcha, MD, MHS,

*Los Angeles Biomedical Research Institute at Harbor UCLA Medical Center, Torrance, CA:* Richard Casaburi, PhD, MD, Alessandra Adami, PhD, Janos Porszasz, MD, PhD, Hans Fischer, MD, PhD, Matthew Budoff, MD, Harry Rossiter, PhD

*Michael E. DeBakey VAMC, Houston, TX:* Amir Sharafkhaneh, MD, PhD, Charlie Lan, DO

*Minneapolis VA:* Christine Wendt, MD, Brian Bell, MD

*Morehouse School of Medicine, Atlanta, GA:* Marilyn Foreman, MD, MS, Gloria Westney, MD, MS, Eugene Berkowitz, MD, PhD

*National Jewish Health, Denver, CO:* Russell Bowler, MD, PhD, David Lynch, MD

*Reliant Medical Group, Worcester, MA:* Richard Rosiello, MD, David Pace, MD

*Temple University, Philadelphia, PA:* Gerard Criner, MD, David Ciccolella, MD, Francis Cordova, MD, Chandra Dass, MD, Gilbert D’Alonzo, DO, Parag Desai, MD, Michael Jacobs, PharmD, Steven Kelsen, MD, PhD, Victor Kim, MD, A. James Mamary, MD, Nathaniel Marchetti, DO, Aditi Satti, MD, Kartik Shenoy, MD, Robert M. Steiner, MD, Alex Swift, MD, Irene Swift, MD, Maria Elena Vega-Sanchez, MD

*University of Alabama, Birmingham, AL:* Mark Dransfield, MD, William Bailey, MD, J. Michael Wells, MD, Surya Bhatt, MD, Hrudaya Nath, MD

*University of California, San Diego, CA:* Joe Ramsdell, MD, Paul Friedman, MD, Xavier Soler, MD, PhD, Andrew Yen, MD

*University of Iowa, Iowa City, IA:* Alejandro Cornellas, MD, John Newell, Jr., MD, Brad Thompson, MD

*University of Michigan, Ann Arbor, MI:* MeiLan Han, MD, Ella Kazerooni, MD, Carlos Martinez, MD

*University of Minnesota, Minneapolis, MN:* Joanne Billings, MD, Tadashi Allen, MD *University of Pittsburgh, Pittsburgh, PA:* Frank Sciurba, MD, Divay Chandra, MD, MSc, Joel Weissfeld, MD, MPH, Carl Fuhrman, MD, Jessica Bon, MD

*University of Texas Health Science Center at San Antonio, San Antonio, TX:* Antonio Anzueto, MD, Sandra Adams, MD, Diego Maselli-Caceres, MD, Mario E. Ruiz, MD

